# Comparison of silhouette-based reallocation methods for vegetation classification

**DOI:** 10.1101/630384

**Authors:** Attila Lengyel, David W. Roberts, Zoltán Botta-Dukát

**Affiliations:** Centre for Ecological Research, Institute of Ecology and Botany, Alkotmány u. 2-4. H-2163 Vácrátót, Hungary; Swiss Federal Research Institute WSL, CH-8903 Birmensdorf, Switzerland; Ecology Department, Montana State University, Bozeman, Montana, USA, 59717-3460

**Keywords:** Flexible-beta, classification, clustering, iterative, OPTIMCLASS, optimization, OPTSIL, REMOS, silhouette, validation

## Abstract

**Aims:** To introduce REMOS, a new iterative reallocation method (with two variants) for vegetation classification, and to compare its performance with OPTSIL. We test (1) how effectively REMOS and OPTSIL maximize mean silhouette width and minimize the number of negative silhouette widths when run on classifications with different structure; (2) how these three methods differ in runtime with different sample sizes; and (3) if classifications by the three reallocation methods differ in the number of diagnostic species, a surrogate for interpretability.

**Study area:** Simulation; example data sets from grasslands in Hungary and forests in Wyoming and Utah, USA.

**Methods:** We classified random subsets of simulated data with the flexible-beta algorithm for different values of beta. These classifications were subsequently optimized by REMOS and OPTSIL and compared for mean silhouette widths and proportion of negative silhouette widths. Then, we classified three vegetation data sets of different sizes from two to ten clusters, optimized them with the reallocation methods, and compared their runtimes, mean silhouette widths, numbers of negative silhouette widths, and the number of diagnostic species.

**Results:** In terms of mean silhouette width, OPTSIL performed the best when the initial classifications already had high mean silhouette width. REMOS algorithms had slightly lower mean silhouette width than what was maximally achievable with OPTSIL but their efficiency was consistent across different initial classifications; thus REMOS was significantly superior to OPTSIL when the initial classification had low mean silhouette width. REMOS resulted in zero or a negligible number of negative silhouette widths across all classifications. OPTSIL performed similarly when the initial classification was effective but could not reach as low proportion of misclassified objects when the initial classification was inefficient. REMOS algorithms were typically more than an order of magnitude faster to calculate than OPTSIL. There was no clear difference between REMOS and OPTSIL in the number of diagnostic species.

**Conclusions:** REMOS algorithms may be preferable to OPTSIL when (1) the primary objective is to reduce or eliminate negative silhouette widths in a classification, (2) the initial classification has low mean silhouette width, or (3) when the time efficiency of the algorithm is important because of the size of the data set or the high number of clusters.

## Introduction

Numerical classification methods are essential data analytical tools in vegetation ecology and several other scientific fields, including genomics, psychology, or sociology. Basically, classification algorithms can be divided into two groups. Hierarchical algorithms produce a perfectly nested hierarchy of clusters of objects, while the output of non-hierarchical methods is a partition in which each classified object is assigned exclusively to one cluster (or, in the special case of fuzzy clustering methods, non-exclusively to several clusters using fuzzy membership weights) at the same level. Hierarchical methods can be subdivided into agglomerative and divisive methods based on whether they initiate the clustering algorithm from treating each single object as a separate cluster, and then merge them until all objects are included in a single cluster at the highest hierarchical level, or they proceed in the opposite direction by dividing the entire sample iteratively into smaller and smaller subsets in a nested way. The diversity of numerical classification methods is reviewed by several authors, e.g. Kaufman & Rousseeuw (1990), Podani (2000), Peet & Roberts (2013), Legendre & Legendre (2012).

The advantage of hierarchical methods is that they do not need a pre-defined cluster number; however, if a single-level classification is the objective, as is generally the case, a hierarchical classification requires a post-hoc assessment for choosing the ‘best’ number of clusters. Moreover, a disadvantage of hierarchical methods is that earlier steps (either merging or division) constrain further ones, hence the final solution may be suboptimal. In such a case the *a posteriori* reallocation of misclassified objects might be necessary.

Recently Roberts (2015) introduced two reallocation-based methods which can be used for improving already existing classifications by optimizing a pre-selected goodness-of-clustering criterion. One of these two, called OPTSIL, optimizes the silhouette width which is a widely used index for evaluating classifications and identifying ‘core’ and misclassified objects individually (Rousseeuw 1987, Kaufman & Rousseeuw 1997). Let *i* be a focal object belonging to cluster *A*. Let *C* be a cluster not containing *i*. *a*(*i*) is defined as the average dissimilarity between *i* and all other objects in *A*, while *c*(*i,C*) is the average dissimilarity between *i* and all objects in *C*.

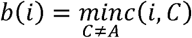

That is, *b(i)* is the average dissimilarity between *i* and the members of its closest neighbour cluster. The silhouette width, *S(i)*, is defined as:

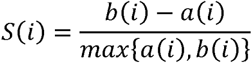

*S(i)* ranges between −1 and +1. Values near +1 indicate that object *i* is much closer to other objects in its assigned cluster than to objects of the closest other cluster, implying a correct classification. If *S(i)* is near 0, the correct classification of the focal object is doubtful, thus suggesting intermediate position between two clusters. *S(i)* values < 0 indicate poor fit, and such objects are often considered ‘misclassified’ (Rousseeuw 1987). In each iteration, OPTSIL evaluates how much the reallocation of any single object in the classification increases the sample-wise mean of silhouette width. It is done by re-assigning each object from its current cluster to every other cluster, and then re-calculating the silhouette widths for all objects. The reallocation which causes the highest increase in the sample-wise mean silhouette is accepted in each step, until no further improvement is possible. Roberts (2015) concluded that OPTSIL is able to significantly improve the initial classification; however, it is slow to converge, and thus recommended for ‘polishing’ of classifications made by other methods.

We present two new silhouette-based reallocation algorithms, called REMOS (reallocation of Misclassified Objects based on Silhouette width). Using artificial and real data sets, we compare them with OPTSIL in terms of three criteria: optimization success, time efficiency, and interpretability.

## Materials and Methods

### The REMOS algorithms

Instead of evaluating the effect of the reallocation of each object (typically sample unit) on the mean silhouette width, REMOS algorithms simply reallocate one or all of the objects which have negative silhouette width. According to how objects to reallocate are selected, we introduce two versions of REMOS. REMOS1 reallocates only the object with the most negative silhouette width (i.e., the ‘worst classified’ object), while REMOS2 reallocates all objects with negative silhouette width (i.e., all misclassified objects). Both algorithms stop if the lowest silhouette width reaches a threshold *L*, or if no further improvement is possible. By default *L* is 0; however, using different values between −1 and 0 can control tolerance towards misclassifications. The steps of the algorithms are presented below:

1. Calculating the silhouette widths, *S(i)*, for the classified objects;
2. Are there any objects with *S(i)* < *L*?
  a. If no, then go to (5)
  b. If yes, go to (3)
3. Updating the classification by reallocating objects: REMOS1: reallocate only the object with the most negative silhouette width to its neighbour cluster; REMOS2: reallocate all the objects with *S(i)* < *L* to their respective neighbour clusters;
4. Go to (1).
5. End – no further optimization is possible

Our preliminary runs showed that both REMOS algorithms frequently converge into loops where the iteration proceeds repeatedly over a finite number of suboptimal solutions without finding any of them as a final solution. To break such a loop, the algorithm checks for repetitions and stops if two identical solutions occur. In this case the solution with the lowest number of negative silhouette widths is selected from the previous iterations. In case of tied minimum of negative silhouette widths, the solution giving the higher absolute sum of negative silhouette widths (a surrogate for smaller ‘classification error’) is chosen as final. Not surprisingly, in most cases REMOS1 requires many more iterations than REMOS2. According to our pilot analyses with differently sized data matrices and different initial classifications, this can extend the computation time of REMOS1 in comparison with REMOS2. It is possible to set an upper limit to the number of iterations; however, as there is no standard value for this threshold, the default setting is infinity (that is, no limit).

An R script of the REMOS algorithms is provided in the Electronic Supplement.

### Data sets

We compared the performance of the REMOS1, REMOS2 and the OPTSIL algorithms on three real and one artificial data set. The Shoshone data set is a random subset comprising 150 plots selected from a larger forest inventory database. This data set represents coniferous forests of Shoshone National Forests (WY, USA). In the plots vascular species were recorded using an ordinal scale. The Bryce data set was sampled in the Bryce Canyon National Park (UT, USA; Roberts 1992). It includes 160 circular plots of ~404.7 m^2^ (0.1 acre) where the cover of 169 vascular species (except trees) were recorded on ordinal scale. The Grasslands data set is a subset of a larger sample of mesic grasslands of northern Hungary (Lengyel et al. 2016). The size of the matrix is 55 plots by 269 species. Abundances are coded on a percentage scale. As artificial data, we employed a simulated data set of 400 points in two dimensions. The points are aggregated into eight fuzzy clusters (Fig. 1). For different test scenarios, random subsets of different size were used.

**Fig. 1.**
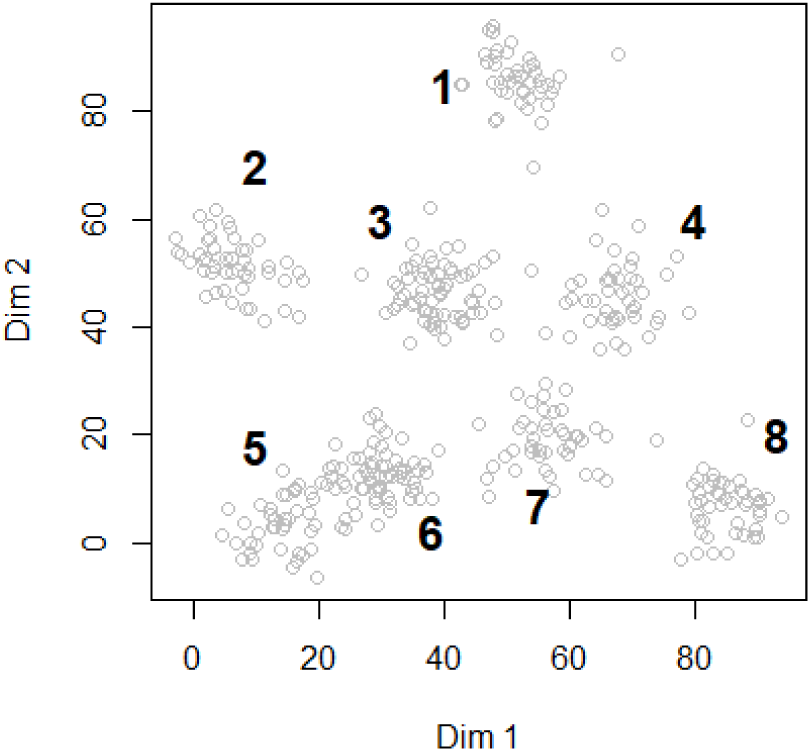
The simulated data set containing 400 points in eight aggregations

### Data analysis

The performance of the REMOS and OPTSIL algorithms was evaluated from three aspects: optimization success on different initial classifications of artificial and real data, dependence of computation time on sample size with artificial data, and interpretability of the optimized classification of real data based on indicator species.

For testing optimization success, initial classifications of random subsamples of the artificial data set containing 200 points were prepared using the flexible-beta classification algorithm (Lance & Williams 1966). This method uses a parameter called beta which enables producing classifications with different sensitivity of ‘chaining’ vs. ‘grouping’ effect. The beta is adjustable between −1 and +1. With lower values the grouping effect is emphasized, while higher beta gives more weight to chaining. With beta = −1 flexible-beta clustering is identical with the complete linkage method, with beta = 0 it agrees with the average linkage (UPGMA), with beta = +1 it is the same as single linkage. Several authors reported that the flexible clustering method provides the most satisfactory classifications using beta = −0.25. In this analysis, values of beta were changed between −1 and +1 in steps by 0.25 in between. The hierarchical classifications were cut at the 8-cluster level. The procedure was repeated 5 times resulting in 5 × 9 = 45 initial classifications. Each of them was optimized using the REMOS1, REMOS2, and OPTSIL algorithms. We compared the change of mean silhouette widths (MSW) and misclassification rate (that is, the proportion of negative silhouette widths; MR) across beta values between the optimized classifications and the initial classification. In the Electronic Supplement we show some exemplary classifications.

For comparing time efficiency, we drew subsamples containing 50, 100, 200, and 300 points of the artificial data set in 20 repeats, and additionally, used also the entire sample of 400 points. Each of them were classified to 8 clusters using the flexible-beta algorithm with beta =-0.25, resulting in 81 initial classifications. These were optimized using REMOS1, REMOS2, and OPTSIL, and the time elapsed during the optimization process was compared between the three algorithms.

Real data sets were classified to 2 to 20 clusters using the flexible-beta algorithm (Lance & Williams 1966) where beta = −0.25. For all real data sets and both classifications, the dissimilarity measure was Sörensen index. Each partition was optimized using the REMOS1, REMOS2, and OPTSIL methods. To assess differences in optimization success, mean silhouette width and misclassification rate were calculated and compared between reallocation methods, the original classification, and across numbers of clusters.

Lötter et al. (2013) argued that species fidelity should be a leading criterion in the evaluation of vegetation classifications. Therefore, we used the Optimclass 1 index as a proxy for interpretability of classifications (Tichý et al. 2010) that is the total number of faithful species across all clusters. Faithful species were determined using Fisher’s exact test and a *p*=0.001 threshold for supporting the null hypothesis that the species shows random distribution across clusters (Chytrý et al. 2002). Hence, we also compared flexible-beta classifications optimized by REMOS1, REMOS2, and OPTSIL, as well as the initial classifications in terms of the number of faithful species across number of clusters.

The data analysis was carried out in the R software environment (R Core Team 2017) using the cluster (Maechler et al. 2018) package. Source code for REMOS1 and REMOS2 is supplied in the Electronic Supplement S3. OPTSIL was calculated using the optpart package (Roberts 2016).

## Results

When comparing optimization success, the REMOS and OPTSIL algorithms differed markedly in MSW values they reached at different values for beta (Fig. 2). With beta <= 0 the mean silhouette width of the initial classification was already high (MSW > 0.60), yet all three optimization methods achieved minor improvement. Within this range of beta, the largest increment in MSW was made by OPTSIL (+0.0134), less by REMOS2 (+0.0105) and REMOS1 (+0.0118) (Table 1). On average, OPTSIL was superior to all other methods in this respect, although differences were very slight (|0.0013| to |0.0029|) between OPTSIL, REMOS1, and REMOS2. From beta = 0.25 and higher, initial classifications showed a dramatic decline in MSW; with beta = 0.5 and higher, MSW dropped below 0. OPTSIL was able to optimize these initial classifications only to a limited degree: MSW ranged between 0.45 and 0.69 with beta = 0.25, and between 0.12 and 0.45 with higher beta. On the contrary, REMOS1 and REMOS2 performed well, achieving a lowest median MSW of 0.599 with beta = 0.75; even the minima were near 0.5. A very similar pattern was detectable with misclassification rates. MR was near 0 with beta <= 0 for both optimization methods (Fig. 3). Within this range, REMOS1 reached the lowest MR on average but its advantage over REMOS2 was minimal (|0.0002| difference; Table 2). REMOS1 and REMOS2 had slightly lower MR than OPTSIL (|0.0034| and |0.0032| differences, respectively). All optimization methods decreased MR in comparison with the initial classification (REMOS1: −0.0187, REMOS2: −0.0185, OPTSIL: −0.0153). With increasing beta, especially with beta >= 0.5, REMOS1 and REMOS2 kept MR at the same level, while OPTSIL resulted in gradually higher values reaching medians over 0.1.

**Table 1.** Differences between the (unoptimized) initial classification, REMOS1, REMOS2, and OPTSIL solutions when the beta = 0 or lower in the flexible-beta classification. In the cells are averages of differences calculated for each run as [MSW by the method in the row] – [MSW by the method in the column].

|  | **Initial** | **REMOS1** | **REMOS2** | **OPTSIL** |
| --- | --- | --- | --- | --- |
| **Initial** | 0 | -0.0105 | -0.0118 | -0.0134 |
| **REMOS1** | 0.0105 | 0 | -0.0013 | -0.0029 |
| **REMOS2** | 0.0118 | 0.0013 | 0 | -0.0016 |
| **OPTSIL** | 0.0134 | 0.0029 | 0.0016 | 0 |

**Table 2.** Differences between the (unoptimized) initial classification, REMOS1, REMOS2, and OPTSIL solutions when the beta = 0 or lower in the flexible-beta classification. In the cells are averages of differences calculated for each run as [MR by the method in the row] – [MR by the method in the column].

|  | **Initial** | **REMOS1** | **REMOS2** | **OPTSIL** |
| --- | --- | --- | --- | --- |
| **Initial** | 0 | 0.0187 | 0.0185 | 0.0153 |
| **REMOS1** | -0.0187 | 0 | -0.0002 | -0.0034 |
| **REMOS2** | -0.0185 | 0.0002 | 0 | -0.0032 |
| **OPTSIL** | -0.0153 | 0.0034 | 0.0032 | 0 |

**Fig. 2.**
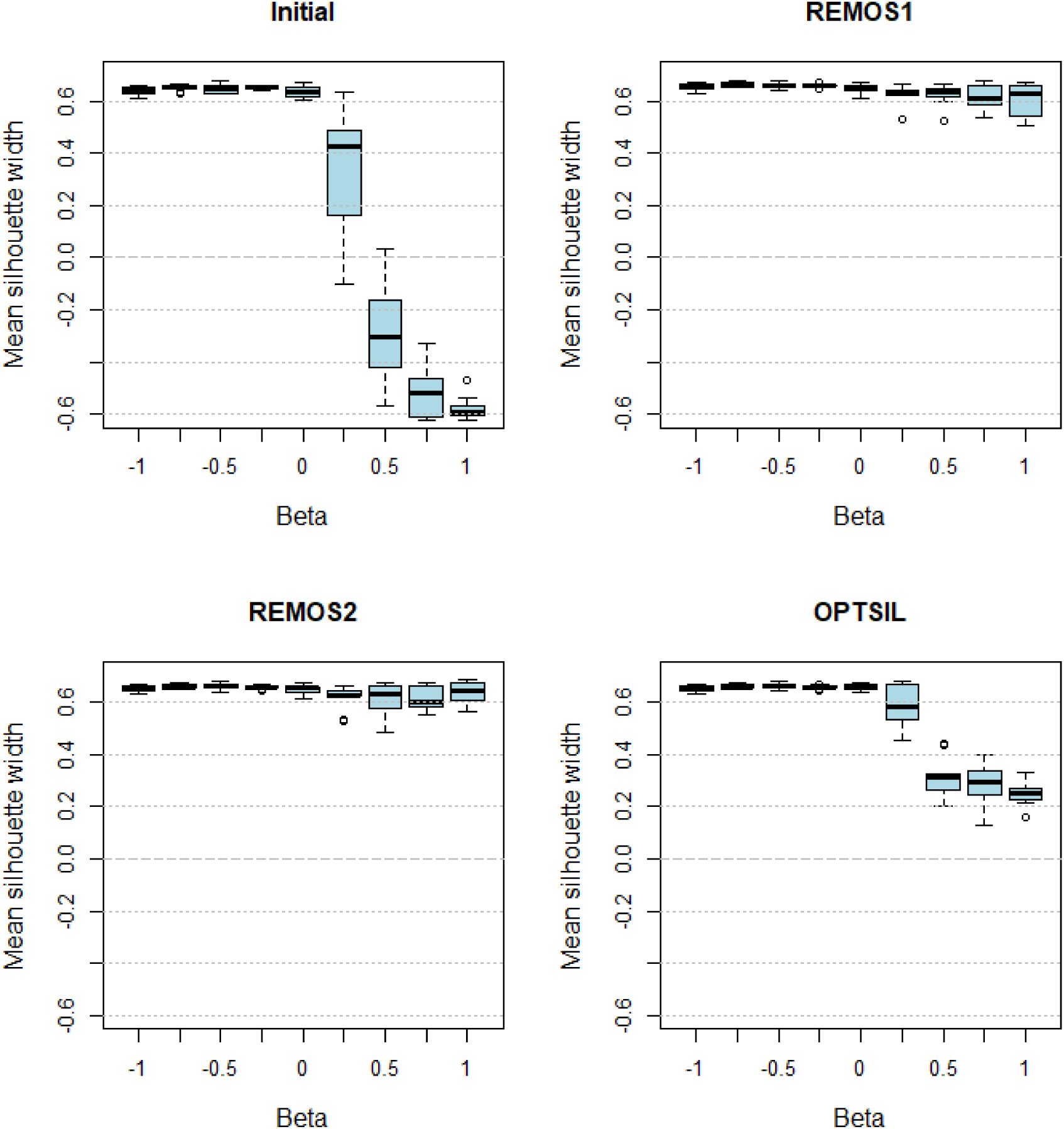
Comparison of the initial classification (without optimization), REMOS1, REMOS2, and OPTSIL across different beta values of the flexible-beta classification based on mean silhouette width.

**Fig. 3.**
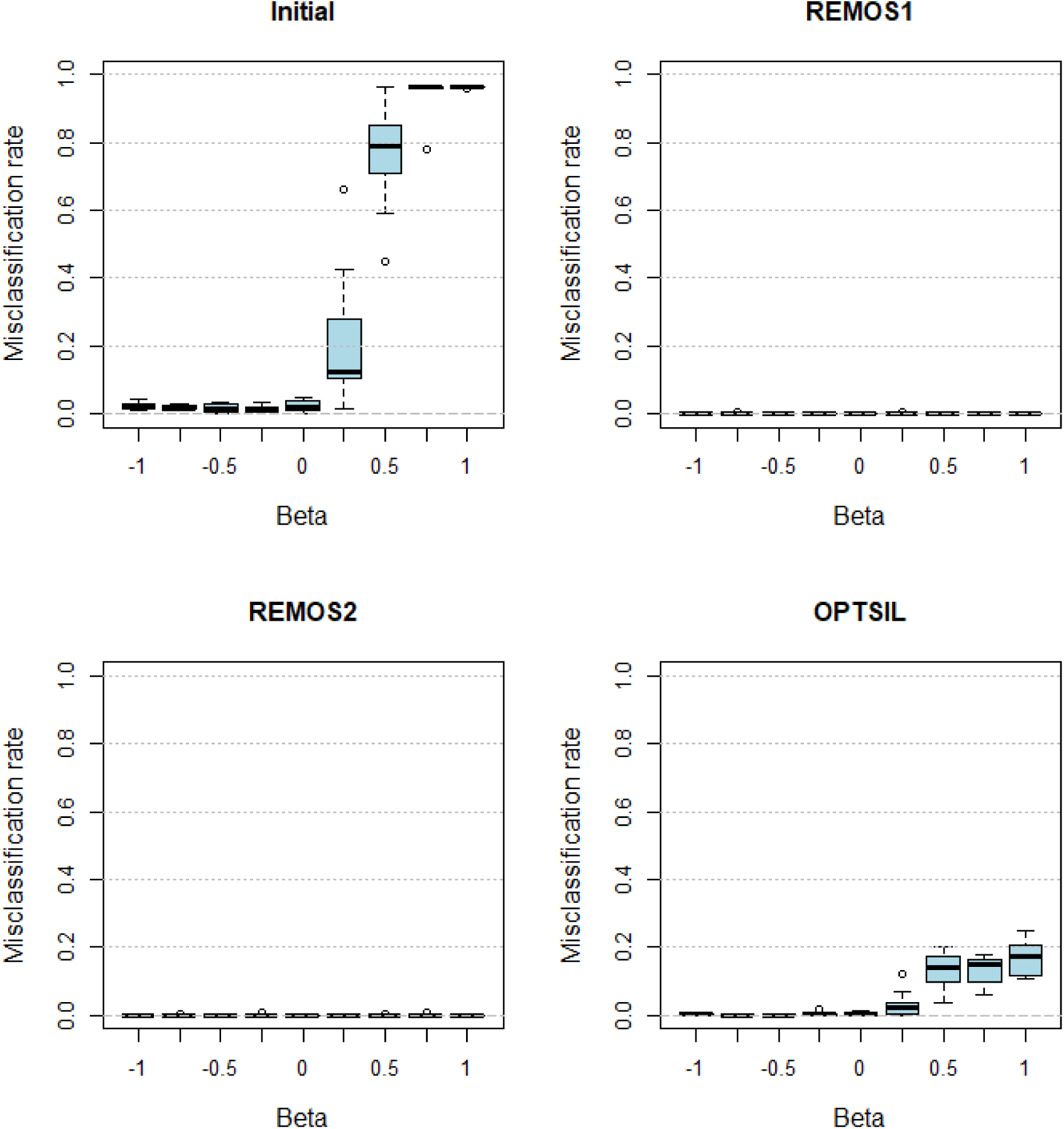
Comparison of the initial classification (without optimization), REMOS1, REMOS2, and OPTSIL across different beta values of the flexible-beta classification based on misclassification rate.

The number of iterations for REMOS1 were between 2 and 234, for REMOS2 between 2 and 53, and for OPTSIL between 0 and 67. Not surprisingly, from less efficient initial classifications more iterations were necessary to reach a final solution; however, the upper limit of number of iterations was never reached.

Visual checking of the classifications showed that with beta = 0 or lower all classifications mirrored the a priori point aggregations efficiently (Figures S4-1 to S4-4). Classifications differed mostly in the assignments of transitional points. With beta > 0 initial classifications tended not to distinguish point aggregations as separate clusters. OPTSIL classification tended to delimit one (or a few) heterogeneous clusters including several aggregations of many points in a single cluster and several clusters with very few points distant from each other (see Fig. S4-7). Additionally, OPTSIL tended to eliminate clusters completely, thus often keeping only 2 to 7 clusters from the initial eight (see Fig. S4-5 to S4-7). In a few cases REMOS2 also eliminated one or two clusters but REMOS algorithms were rather consistent in delineating point aggregations rather independently of the beta value.

There was a significant difference in the relationship between sample size and computation time among the three optimization methods (Fig. 4). Considering average runtimes with 50 points REMOS2 was the fastest (0.0006 s), followed closely by REMOS1 (0.0010), while OPTSIL ran approximately three times longer (0.0254 s). For larger samples, REMOS2 was the fastest, completing classifications in 0.0008 s, 0.0010 s, 0.0018 s, and 0.0010 s, with 100, 200, 300, and 400 objects, respectively. However, its advance over REMOS1 was minimal, which needed 0.0015 s, 0.0025 s, 0.0051 s, 0.0030 s on average. The lag of OPTSIL was even more significant at these sample sizes: average runtimes were 0.2556 s, 3.7027 s, 13.7790 s, and 24.4539 s with 100, 200, 300, and 400 objects.

**Fig. 4.**
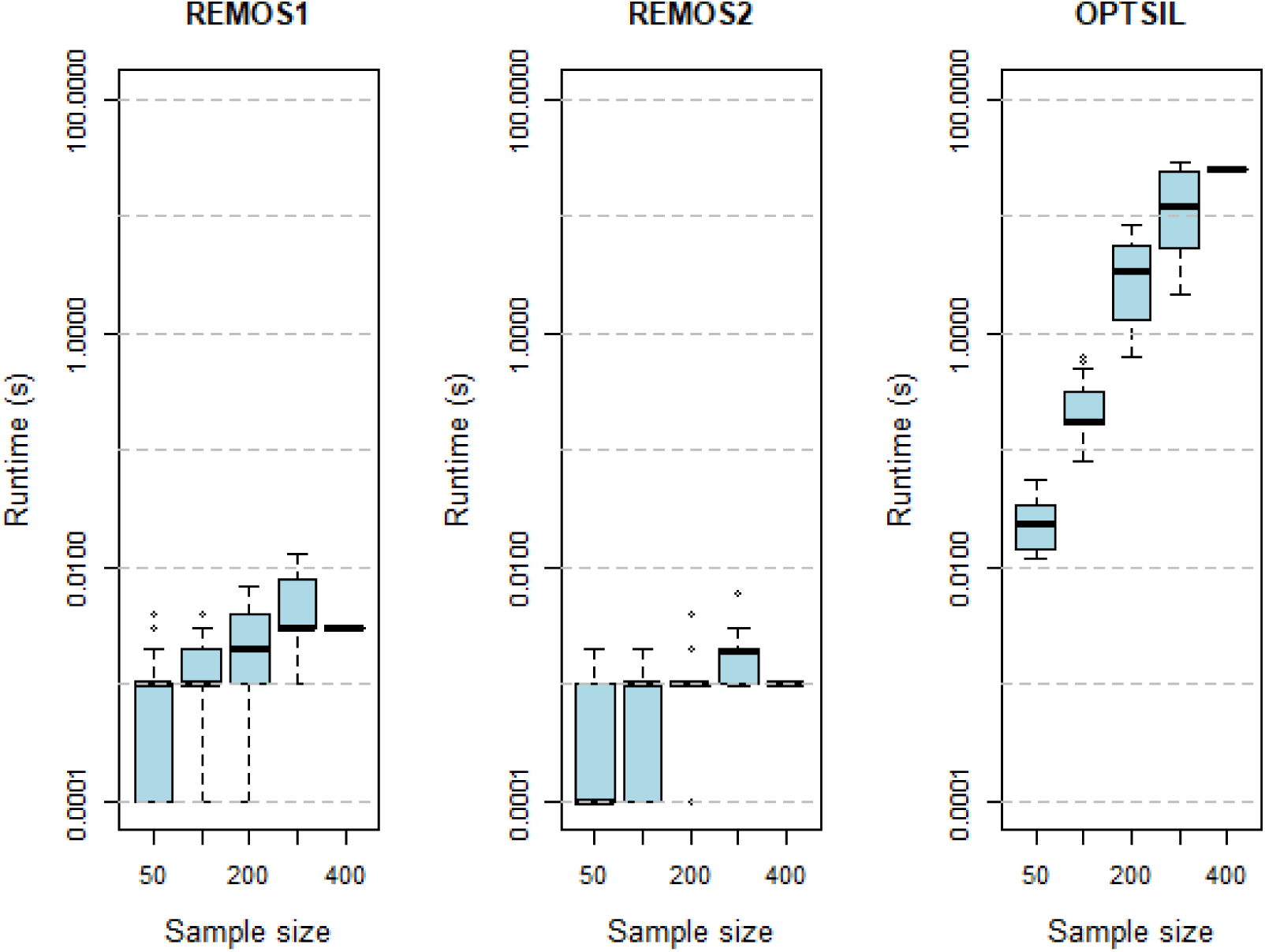
Computation times with different sample sizes by REMOS1, REMOS2, and OPTSIL. Shortest computation times are truncated and replaced by 0.0001 s.

On the Grasslands data set OPTSIL reached the highest MSW at all but two examined cluster levels (Fig. 5). With 6 and 10 clusters REMOS1 performed the best and it was only slightly worse than OPTSIL in all other cases. Interestingly, REMOS2 gave the same MSW values with 2 to 5 clusters (likely due to identical final solutions), but at finer resolutions it was much poorer. With 6, 7 and 9 clusters REMOS2 even decreased the MSW of the initial classification. Regarding misclassification rate, REMOS1 performed the best with no negative silhouette width values over all runs. As with MSW, from 2 to 5 clusters REMOS2 gave the same result, but the weak performance with 6 or more clusters was visible here, too. OPTSIL solutions ranked in intermediate position between REMOS1 and the initial classification, the latter being the worst in all but two cases. These differences were not observable with diagnostic species. Between 2 to 7 clusters all methods (including the initial classification) showed similar numbers of diagnostic species, while at finer resolutions REMOS2 was the best. Nevertheless, the Grassland data set is small, thus at this level the sizes of clusters are so small and the number of diagnostic species so low that these differences are probably not relevant.

**Fig. 5.**
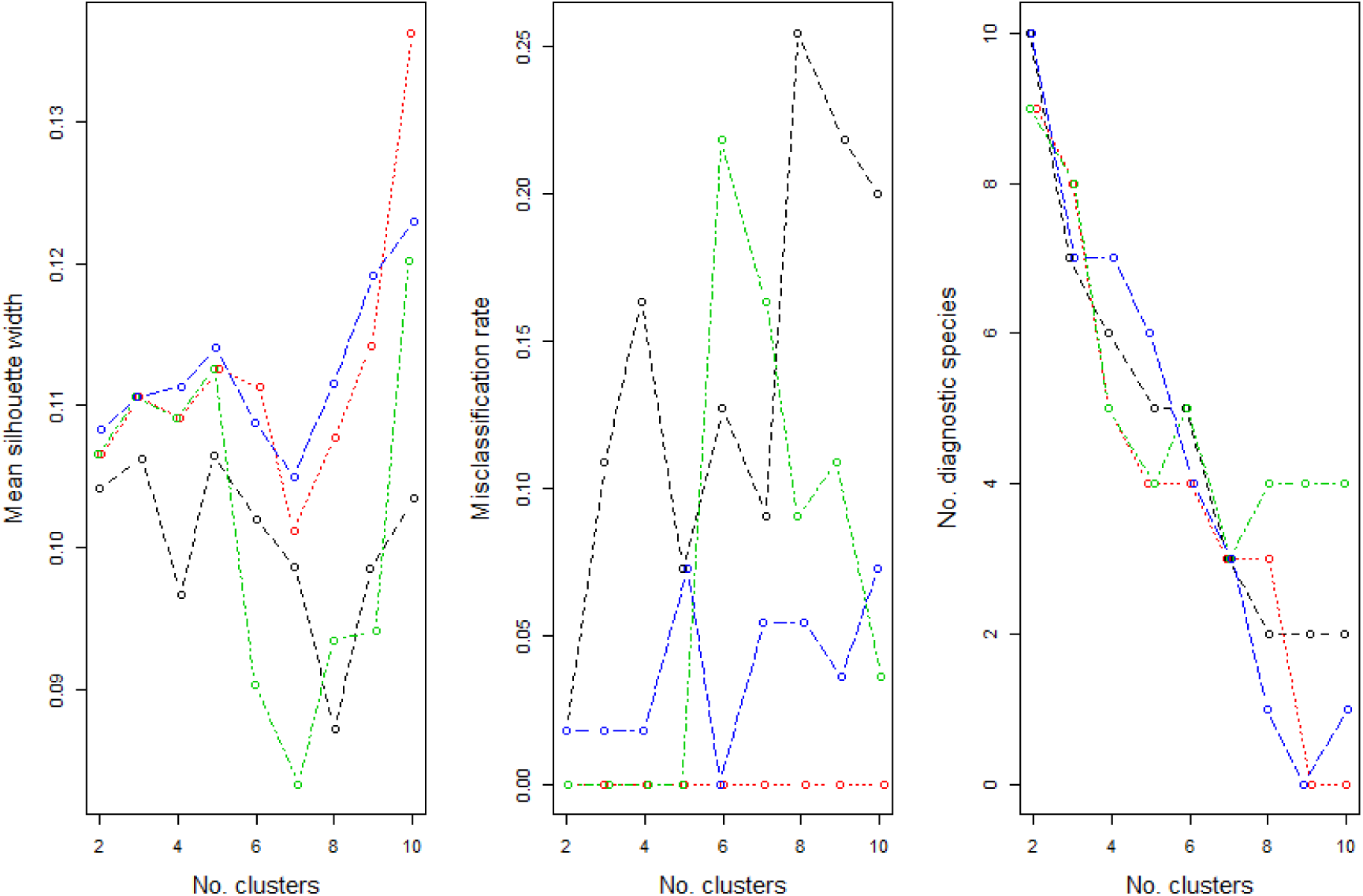
Comparison of the initial classification (without optimization), REMOS1, REMOS2, and OPTSIL solutions in terms of the change of mean silhouette width and number of diagnostic species across the number of clusters on the Grassland data set. The initial classification was produced by the flexible-beta method (beta = −0.25). To avoid overlap, points are jittered in the horizontal direction on the graph. Colour code: red – REMOS1, green – REMOS2, blue – OPTSIL, black – initial classification.

With the Bryce data set OPTSIL produced the highest MSW at most cluster levels (Fig. 5). REMOS1 and REMOS2 had very similar, often identical performance. With a minimal difference they outperformed OPTSIL at two clusters. At 3 and 4 clusters they were slightly worse than OPTSIL but this difference increased with the number of clusters, and became striking from 7 and more clusters. The initial classification had the lowest MSW across the tested numbers of clusters. REMOS1 and REMOS2 provided solutions with the lowest MR, most often with no negative silhouette widths at all. OPTSIL had MR between 0.02 and 0.07, while the initial classification had the highest MR in at all cluster numbers (MR between 0.048 and 0.15). OPTSIL performed the best in terms of diagnostic species at 3, as well as at 6 and more clusters. Interestingly, at 4 clusters the initial classification had the most diagnostic species, while at 2 and 5 clusters REMOS algorithms reached the highest values.

On the Shoshone data set, OPTSIL reached the highest MSW across all cluster numbers, REMOS1 was the second best, showing similar (in a few cases identical) MSW values with REMOS2, and the worst was the initial classification (Fig. 6). REMOS1 had the lowest MR again. This position was shared with REMOS2 between 2 and 5 clusters when both algorithms provided no misclassifications. OPTSIL had MR between 0.04 and 0.07, which positioned it behind REMOS2 in all but two cluster numbers. The initial classification had the highest MR (between 0.15 and 0.27). Regarding the number of diagnostic species, the picture was different. REMOS1 gained the highest numbers, again in a few cases together with REMOS2, while OPTSIL was always inferior. The initial classification was again the worst in all cases, except for the 10-cluster level, where REMOS2 had the fewest diagnostic species.

**Fig. 6.**
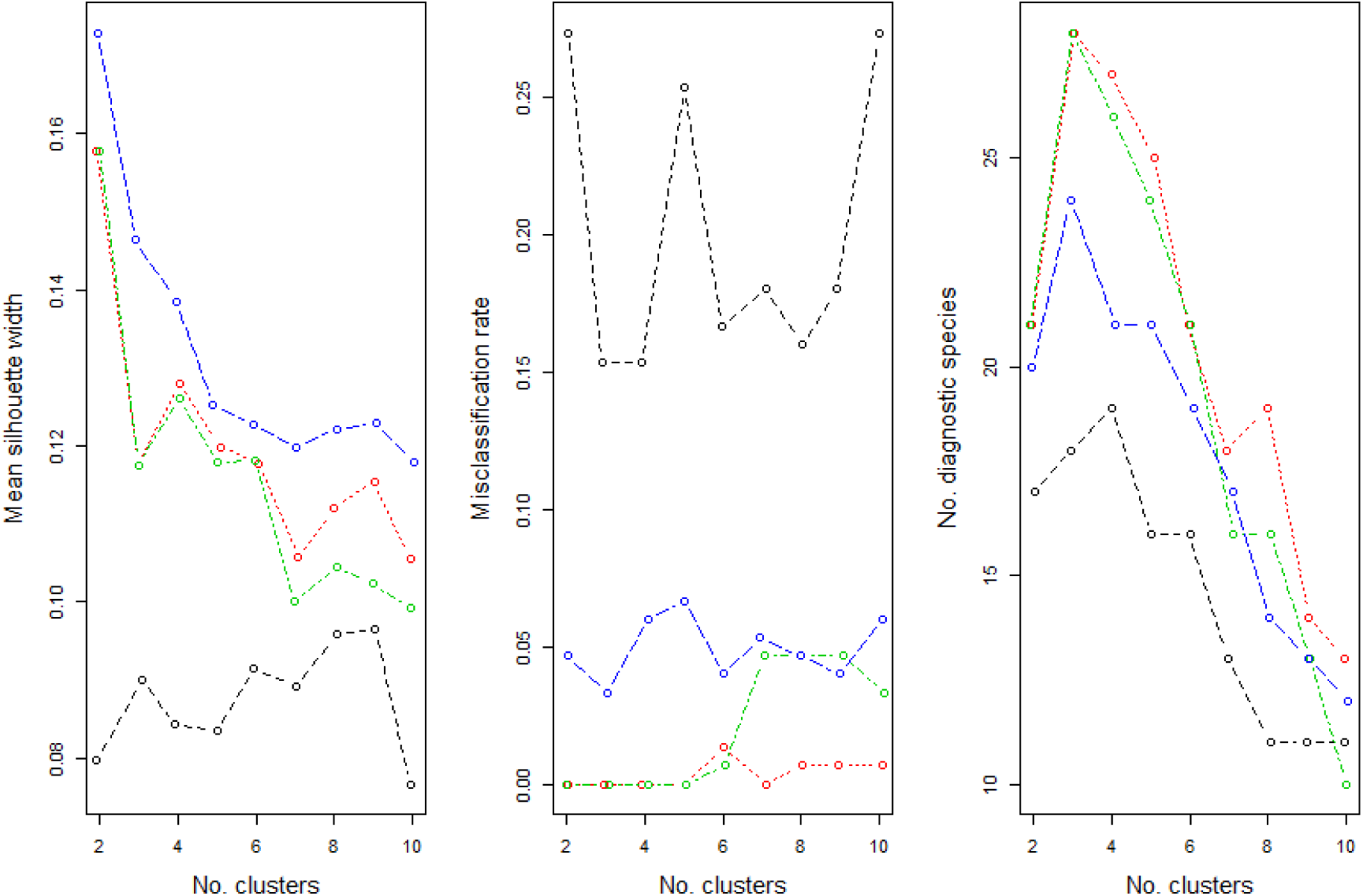
Comparison of the initial (without optimization) classification, REMOS1, REMOS2, and OPTSIL solutions in terms of the change of mean silhouette width and number of diagnostic species across the number of clusters on the Shoshone data set. The initial classification was produced by the flexible-beta method (beta = −0.25). To avoid overlap, points are jittered in horizontal direction on the graph. Colour code: red – REMOS1, green – REMOS2, blue – OPTSIL, black – initial classification.

## Discussion

In this paper we introduced the REMOS algorithms which can be used for improving already existing classifications by reallocating misclassified objects using the silhouette criterion. Two versions are available: REMOS1 reallocates only the single object with the lowest silhouette width, while REMOS2 re-assigns all objects with negative silhouette width to their respective closest neighbour cluster. We provide evidence on the high optimization success and time efficiency of the new algorithms.

Our tests showed that the efficiency of the tested reallocation algorithms (REMOS1, REMOS2 and OPTSIL) has different degrees of dependence on the initial classification. Regarding MSW, the optimization success of OPTSIL is higher than REMOS algorithms’ when the initial classification already has rather high MSW; although, the difference is usually small. Since mean silhouette width prefers spherical cluster shapes (Rousseeuw 1987), it is typically high for classifications produced by group-forming methods, e.g. flexible-beta with beta <= 0, and similar behaviour can be expected when applied to average linkage, complete linkage, Ward’s method, K-means, or PAM classifications. However, with chaining algorithms, e.g. beta > 0, REMOS1 and REMOS2 outperform OPTSIL. Chaining algorithms optimize on criteria emphasising nearest neighbour distances which are not well reflected by the traditional form of silhouette width also applied here (but see Lengyel & Botta-Dukát in press), resulting in non-spherical clusters and low MSW. Using such classifications as input, OPTSIL frequently converges into local optima, while REMOS algorithms provide more robust optimization and reach high MSW. As it was shown by our examples, OPTSIL solutions in these situations often fail to mirror the original cluster structure of the data set. In concurrence with Roberts (2015), we suggest classifying the data set by a grouping method first, and then optimizing the result with OPTSIL in order to reach the highest possible MSW. Alternatively, REMOS algorithms seem more effective with other types of initial classifications, although, their final MSW might be slightly lower than what is maximally possible with OPTSIL.

Regarding misclassification rate, REMOS1 performed the best. In many cases REMOS2 led to exactly the same solution containing no negative silhouette width values at all; however, with the real data examples and higher number of clusters REMOS2 tended not to reach such efficiency. The sensitivity to the initial classification of OPTSIL was visible also on the presence of negative silhouette widths: OPTSIL had significantly higher MR than REMOS algorithms when initiated from classifications with a chained structure. It must be noted that different algorithms may reach the same value for MR, while their final solutions are not necessarily identical. It occurred in some times with REMOS1 and REMOS2 that their final solutions contained no, or only very few misclassified objects, while their classifications were different. Even the number of clusters can differ between REMOS1 and REMOS2 despite equal MR (e.g., Fig. S4-4). Such agreement in MSW is less probable due to its continuous scale.

In general, optimizing a single criterion results in trade-offs for other criteria, and OPTSIL and REMOS demonstrate this clearly. It is not surprising that OPTSIL reached the highest MSW values, while REMOS outperformed OPTSIL in terms of MR. When comparing the optimization success of OPTSIL and REMOS on MSW and MR, it must be noted that OPTSIL directly maximizes MSW, a ‘global’ criterion of classification efficiency. REMOS, on the other hand, has a more local perspective on classification efficiency, and focuses on neighbourhoods of adjacent clusters. Surely, MSW and MR correlate strongly, and in general optimizing MR will lead to high, although not necessarily optimal, MSW. In addition, REMOS implicitly minimizes the absolute value of the sum of negative silhouette widths. In our tests, this criterion behaved very similarly to MR, thus we present its results only in the Electronic Supplement 5.

OPTSIL employs an anticipatory algorithm that tentatively reallocates an object to another cluster, but then calculates the consequences of doing so before making the reallocation effective. As a result, the trace of the optimization criterion is strictly monotonic increasing. REMOS, on the other hand, identifies candidate objects to reallocate and makes the reallocation effective immediately. In some cases this causes objects in the target cluster to exhibit newly negative silhouette widths in the next iteration, and subsequent reallocations must undo the negative consequences of a previous reallocation. As a result, the trace of the optimization criterion shows non-monotonic behaviour, and in some cases oscillates or exhibits cycles. While in general this behaviour is undesirable it may help avoid local optima in a manner similar to genetic algorithms.

The difference between ‘global’ vs ‘local’ perspective can be seen on the classifications of the artificial data sets (see the Electronic Supplement). OPTSIL solutions initiated from less efficient classifications often contained one or more clusters with a single object, or a few objects which were distant from each other (e.g., Figure S4-7). Such solutions are presumed to have the highest possible MSW from the respective initial classification with the cost of a few very heterogeneous or overlapping clusters and misclassified objects.

From the perspective of optimizing silhouette width, it is not correct to say that an object with negative silhouette width is misclassified if reallocating it to its nearest neighbour cluster decreases MSW. Rather, a misclassification is an assignment that lowers mean silhouette width. However, as noted above, MSW cannot be high with many negative silhouette widths. Alternatively, the viewpoint that correct classification reflects strictly positive silhouette widths for as many objects as possible might be more straightforward than an ‘on-average correct’ solution. This requires a decision from the investigator before choosing between these methods.

An important property of REMOS2 and OPTSIL is that they are able to eliminate complete clusters from the initial classification, thus the final number of clusters becomes lower than the initial. This can be useful if the initial classification has more clusters than is optimal. However, our simulation examples showed that the number of clusters by these methods, but especially OPTSIL, can decrease even if the initial classification is not effective, despite its cluster number corresponds to the number of point aggregations.

We found clear difference in computation time between the three methods. REMOS algorithms were magnitudes faster than OPTSIL. This is not surprising considering that in every iteration of the OPTSIL algorithm all possible reallocations of all objects to each cluster are recalculated and only the one bringing the highest increment in MSW is accepted. In our tests for computation time we used rather small data sets (i.e., containing max. 400 objects) with clear cluster structure, and optimized initial classifications with relatively high MSW. Presumably, such classifications would be faster to optimize than real data sets. Therefore, our measured runtimes are likely to be shorter than what we can expect for larger and more complicated data sets, less efficient classifications or more clusters. If time efficiency of the analysis is crucial and the small difference in optimization success can be neglected, REMOS1 or REMOS2 should be considered instead of OPTSIL.

Although we found OPTSIL sensitive to the initial classification, we must note that OPTSIL performed poorly in situations which are scarcely realistic, since chaining algorithms are rarely used in practice. If high silhouette width is a desired outcome it makes little sense to begin with a classification emphasizing connectivity (e.g. single linkage or flexible-beta with beta > 0), and classifications emphasizing cluster disjunction (e.g. complete linkage or flexible-beta with beta < 0) should be preferred. In vegetation science, group-forming methods are much more popular and straightforward, thus these drawbacks of OPTSIL may not obtain in practice.

Tests on real data showed that OPTSIL combined with flexible-beta (beta = −0.25) is more efficient than REMOS algorithms in terms of MSW, although, the difference is often small. As a contrast, with respect to minimizing the proportion of negative silhouette widths REMOS1 provided consistently the best classifications. However, these differences may not affect interpretability the same way since we could not detect consistent difference between OPTSIL and REMOS algorithms in the number of diagnostic species. We suggest considering which cluster validity measure fits the research question the best, and then decide between the methods discussed above.

## Conclusions

We present REMOS1 and REMOS2 as new reallocation methods for the optimization of classifications and compare them with the related OPTSIL algorithm. When the initial classification is already relatively efficient, most frequently OPTSIL gives the highest final mean silhouette width; however, REMOS solutions are often only slightly worse. When the initial classification has low mean silhouette width, OPTSIL performs poorly, while REMOS algorithms are similarly straightforward as with more efficient initial classifications. With respect to the proportion of misclassified objects, REMOS algorithms, especially REMOS1, provided better classifications than OPTSIL, and this difference increased toward less efficient initial classifications. REMOS algorithms are much time efficient to compute than OPTSIL. We found no systematic difference in the number of diagnostic species between vegetation classifications obtained by OPTSIL and REMOS algorithms.

## Supporting information

R code for the REMOS algorithms

R code for simulated data

Grassland data set

Exemplary classifications of the simulated data set

Comparison of reallocation methods on real data sets using misclassification rate and the absolute sum of negative silhouette widths

## Author contributions

A.L. raised the idea, wrote the scripts, did data analysis, lead writing; D.W.R. did data analysis, discussed results, contributed to the manuscript; Z.B.D. discussed results, commented on the manuscript.

## Data availability

The simulated and the Grassland data sets are available in the attachment. Bryce and Shoshone data sets are available through the labdsv and optpart R packages respectively.

## Acknowledgements

The work of A.L. was supported by the National Research, Development and Innovation Office, Hungary (project number PD-123997).

## Abbreviations

MSW: mean silhouette width
MR: misclassification rate

## Electronic Supplements

Supplement S1: the R code of REMOS

Supplement S2: the R code for the simulated data set

Supplement S3: the Grassland data set (in txt format directly readable to R) Supplement S4: Exemplary classifications of the simulated data set

Supplement S5: Comparison of reallocation methods on real data sets using misclassification rate and the absolute sum of negative silhouette widths

