## Supplementary material for "Comparison of silhouette-based reallocation methods for vegetation classification": R code for the REMOS algorithms

Electronic Supplement for the paper

**Supplement S1 – R code for the REMOS algorithms**

```
remos<-function(d,gr,lim=-0.001,method=c(1,2), return.partitions=F, max.iter=Inf)  {
  gr_new<-gr
  o<- -1
  z<-1
  dw<-1
  w<-u<-y<-vector('numeric')
  W<-matrix(NA, nrow=length(gr), ncol=0)
  parts<-vector('numeric')
  while(o<lim)  {
    SIL<-cluster::silhouette(gr_new,d)
    orig<-SIL[,1]
    neig<-SIL[,2]
    width<-SIL[,3]
    W<-cbind(W,width)
    if(method==1) worst<-which.min(width)
    if(method==2) worst<-which(width<lim)
    if(z>1) dw<-as.vector(colSums((W-width)^2)[-z])
    o<-min(width)
    if(o<lim) {
     gr_new[worst]<-neig[worst]
    }
    y[z]<-o
    u[z]<-sum(width<0)
    w[z]<-abs(sum(width[width<0]))
    parts<-cbind(parts,gr_new)
    if(any(dw==0))   o<-lim+1
    if(z==max.iter) o<-lim+1
    z<-z+1
    }
  fin1<-u==min(u)
  fin2<-w==min(w[fin1]) & fin1
  gr_final<-parts[,fin2]

  if(return.partitions==T) result<-list(gr_final,parts,y,u,w)
  if(return.partitions==F) result<-list(gr_final,y,u,w)
  return(result)
}
```

d: a distance object

gr: an integer vector coding the a priori classification of sample units

lim: a threshold of silhouette width for misclassified objects. It is basically 0 but can be changed to any value between -1 and 0.

method: 1 for REMOS1, 2 for REMOS2.

return.partitions: T for returning intermediate classifications

max.iter: maximal number of iterations

The result is the optimized classification in a form of a vector of integers.

Dependency: cluster package (Maechler et al. 2018).
