## Supplementary material for "Comparison of silhouette-based reallocation methods for vegetation classification": R code for simulated data

Electronic Supplement for the paper

**Supplement S2 – R code for simulated data**

ss<-308

set.seed(ss)

N<-400

K<-8

meanx<-sample(1:100, K)

meany<-sample(1:100, K)

wh<-sample(1:K,N,replace=T)

means<-cbind(meanx,meany)

x<-rnorm(N,mean=meanx[wh],5)

y<-rnorm(N,mean=meany[wh],5)

X<-cbind(x,y)
