## Supplementary material for "Comparison of silhouette-based reallocation methods for vegetation classification": Exemplary classifications of the simulated data set

Electronic Supplement for the paper

**Supplement S4 – Exemplary classifications of the simulated data set**

**Abbreviations:** MSW – mean silhouette width; MR – misclassification rate

**Figure S4-1**. Example – an initial classification with beta flexible method (beta = -1.0) without optimization and with optimization using REMOS1, REMOS2, and OPTSIL. The data set is a random sample containing 200 points drawn from the artificial data set (see main text). A priori point aggregations are numbered. The value of K shows the number of clusters. Delimited clusters are differentiated by colours.

With negative beta, the initial classification used to detect successfully the a priori structure of point aggregations and have high MSW. In this case there was minimal difference between optimization methods. OPTSIL provided slightly higher MSW than REMOS algorithms, while all optimization methods reached the same value for MR. The difference between REMOS and OPTSIL is due to the different delimitation of points in marginal position with respect to their aggregation. In real situation such differences are usually not relevant.

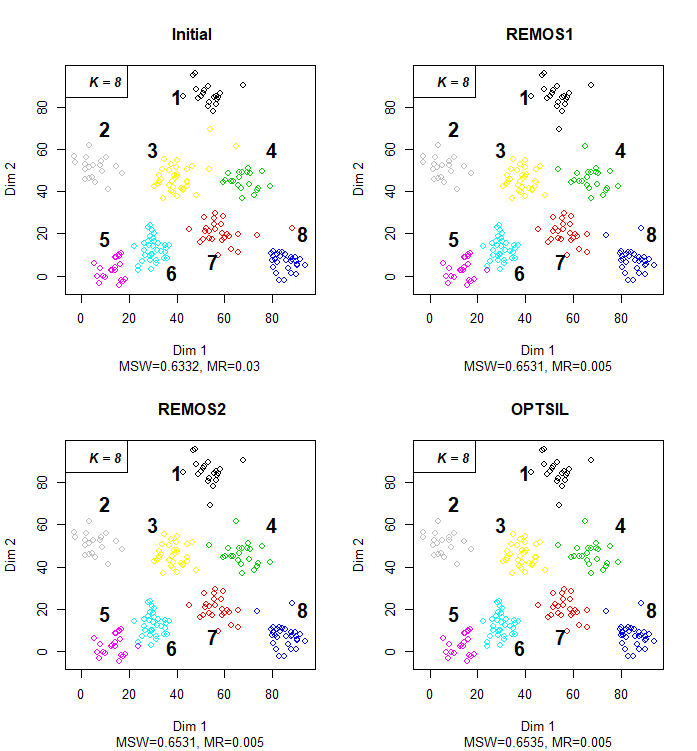

**Figure S4-2**. Initial classification with beta flexible method (beta = -0.5) without optimization and with optimization using REMOS1, REMOS2, and OPTSIL. The data set is a random sample containing 200 points drawn from the artificial data set (see main text). A priori point aggregations are numbered. The value of K shows the number of clusters. Delimited clusters are differentiated by colours.

Another example with negative beta, where initial MSW was high and MR was low. REMOS solutions had slightly higher MSW and lower MR (actually, with no misclassified object), while OPTSIL had the highest MSW, and same low value for MR as the initial classification. The minor differences come from the delimitation between aggregation 5 and 6.

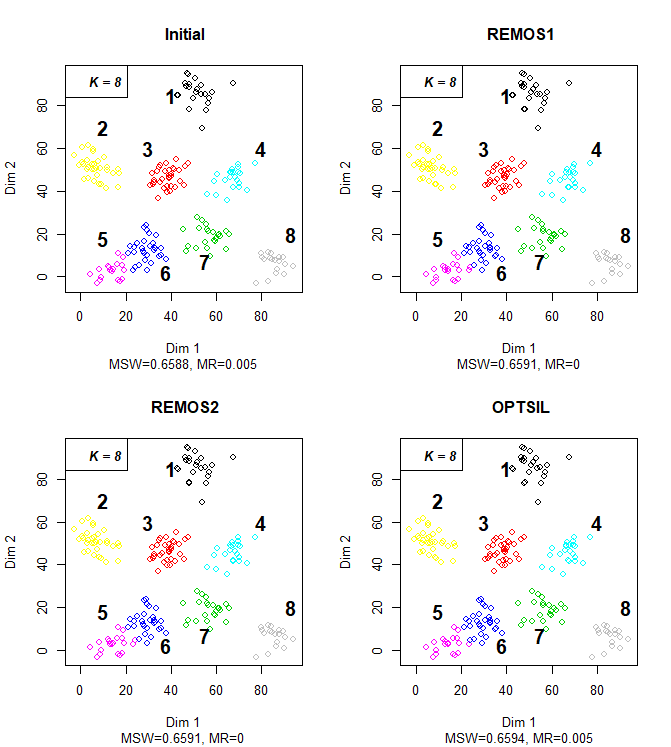

**Figure S4-3**. Initial classification with beta flexible method (beta = -0.25) without optimization and with optimization using REMOS1, REMOS2, and OPTSIL. The data set is a random sample containing 200 points drawn from the artificial data set (see main text). A priori point aggregations are numbered. The value of K shows the number of clusters. Delimited clusters are differentiated by colours.

This example shows a frequent case within our simulations with negative beta, when REMOS algorithms and OPTSIL gave exactly the same result. There was only one point (between aggregates 7 and 8) to relocate in the initial classification to reach the optima for REMOS and OPTSIL.

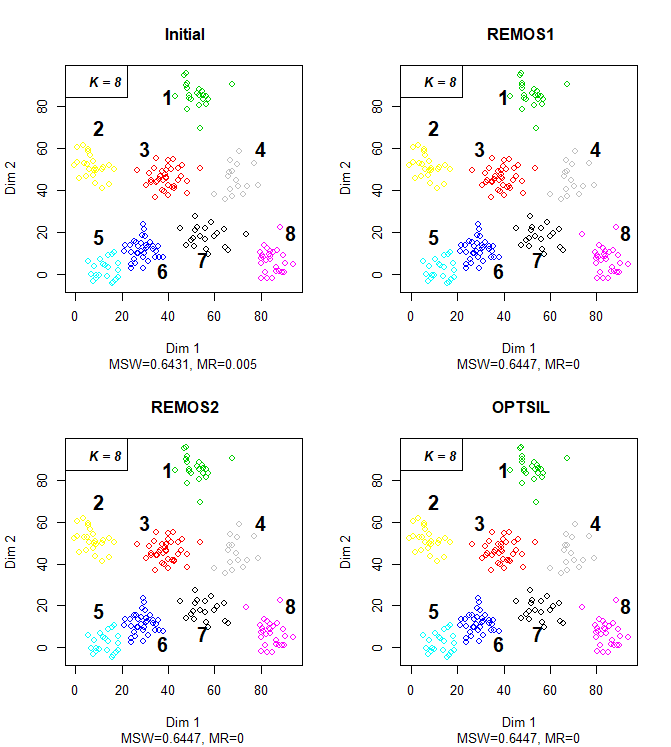

**Figure S4-4**. Initial classification with beta flexible method (beta = 0.0) without optimization and with optimization using REMOS1, REMOS2, and OPTSIL. The data set is a random sample containing 200 points drawn from the artificial data set (see main text). A priori point aggregations are numbered. The value of K shows the number of clusters. Delimited clusters are differentiated by colours.

With beta = 0 MSW and MR values remained in the similar range as with negative beta. In this example, REMOS2 performed the best and OPTSIL reached the second highest MSW. REMOS2 completely eliminated the grey cluster, while OPTSIL assigned two distant objects to it. Interestingly, REMOS1 decreased MSW in comparison with the initial classification by assigning some points of aggregation 1 to the grey cluster. Both REMOS1 and REMOS2 reached MR = 0, while for OPTSIL and the initial classification was MR = 0.01.

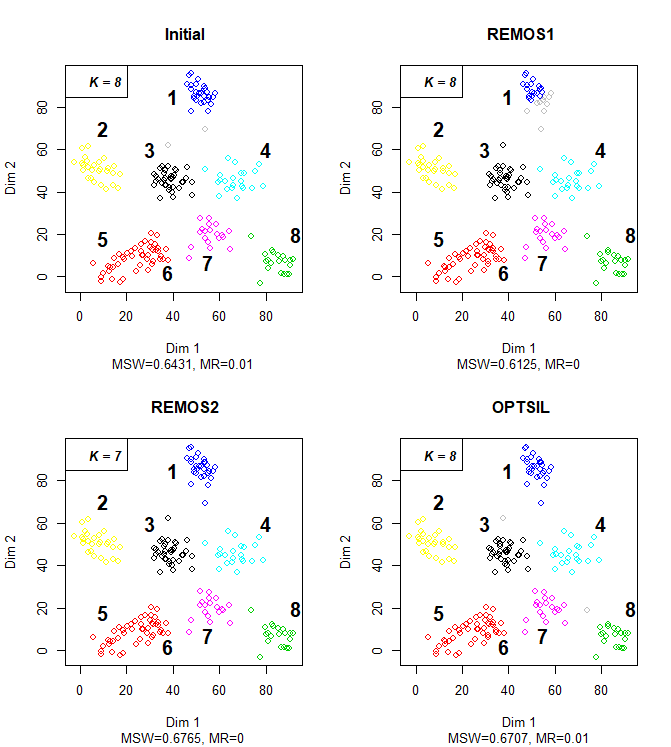

**Figure S4-5**. Initial classification with beta flexible method (beta = 0.25) without optimization and with optimization using REMOS1, REMOS2, and OPTSIL. The data set is a random sample containing 200 points drawn from the artificial data set (see main text). A priori point aggregations are numbered. The value of K shows the number of clusters. Delimited clusters are differentiated by colours.

With positive beta the MSW of initial classification was lower and the MR was higher than in the above examples. With beta = 0.25 REMOS algorithms often reached higher MSW than OPTSIL but not in the example shown below. However, REMOS solutions did not contain misclassified objects, while OPTSIL had a six of them. OPTSIL eliminated the grey cluster and delimited blue and red clusters as similarly heterogeneous as in the initial classification. No method could solve the initial heterogeneity of the blue cluster including two aggregations, 3 and 4.

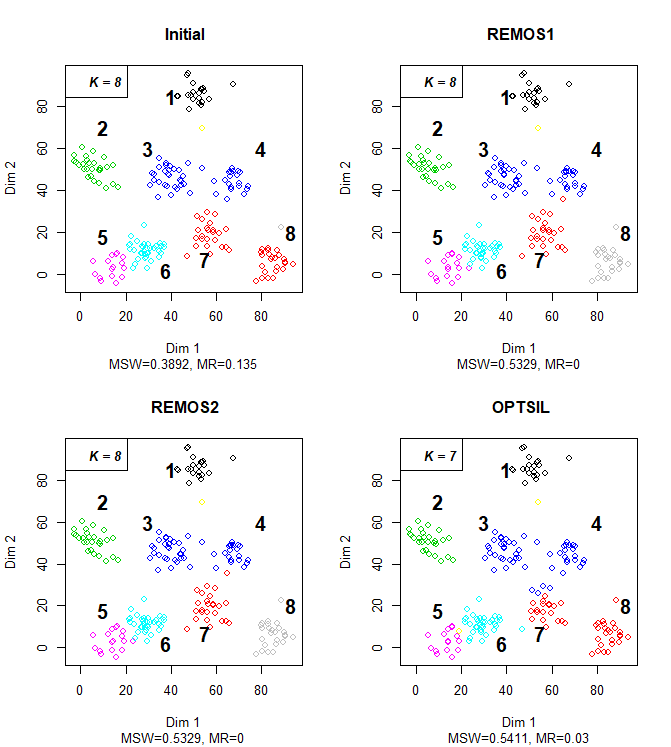

**Figure S4-6**. Initial classification with beta flexible method (beta = 0.5) without optimization and with optimization using REMOS1, REMOS2, and OPTSIL. The data set is a random sample containing 200 points drawn from the artificial data set (see main text). A priori point aggregations are numbered. The value of K shows the number of clusters. Delimited clusters are differentiated by colours.

From beta = 0.5 in the initial classifications clusters often had irregular shape and unequal size, as well as MSW near 0 and high MR. REMOS algorithms were able to reach similar final solutions to those obtained with negative beta but this could not to be said about OPTSIL. OPTSIL delimited only six clusters out of the initial eight. The black and the blue clusters of OPTSIL corresponded well with single aggregations, while the red and the green ones were heterogeneous, including several aggregations. The cyan and the violet clusters were small and widely dispersed. This type of classifications were frequent with beta = 0.5 and higher.

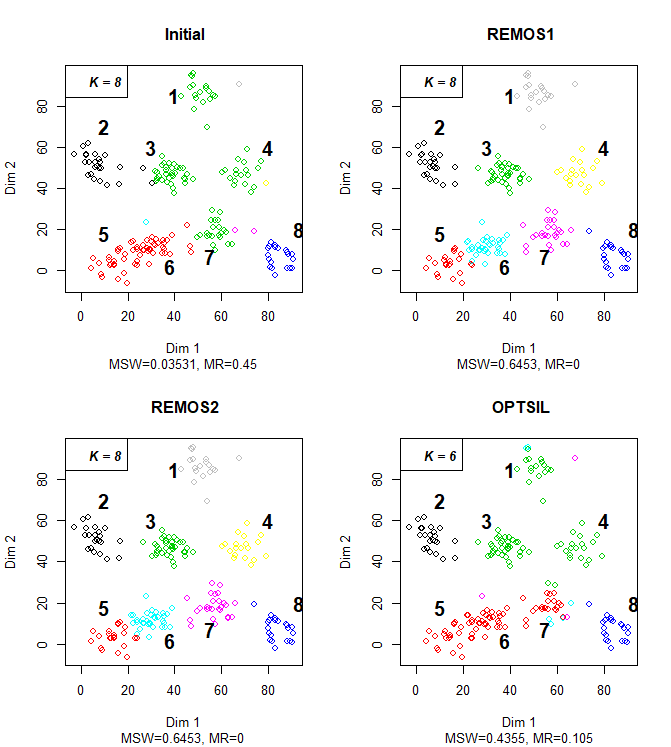

**Figure S4-7**. Initial classification with beta flexible method (beta = 1.0) without optimization and with optimization using REMOS1, REMOS2, and OPTSIL. The data set is a random sample containing 200 points drawn from the artificial data set (see main text). A priori point aggregations are numbered. The value of K shows the number of clusters. Delimited clusters are differentiated by colours.

With beta = 1.0 initial classification mostly comprised of several clusters containing only a single point and the rest points being assigned to a single cluster. As in this example, REMOS algorithms achieved high MSW and MR = 0 even from such initiations, and the final solutions mirrored the original point aggregations very accurately. OPTSIL kept one large cluster (the blue one) containing several point aggregations, while aggregation 8 was divided between three overlapping clusters (black, red, green). Interestingly, all these three clusters had one point also from aggregation 5.

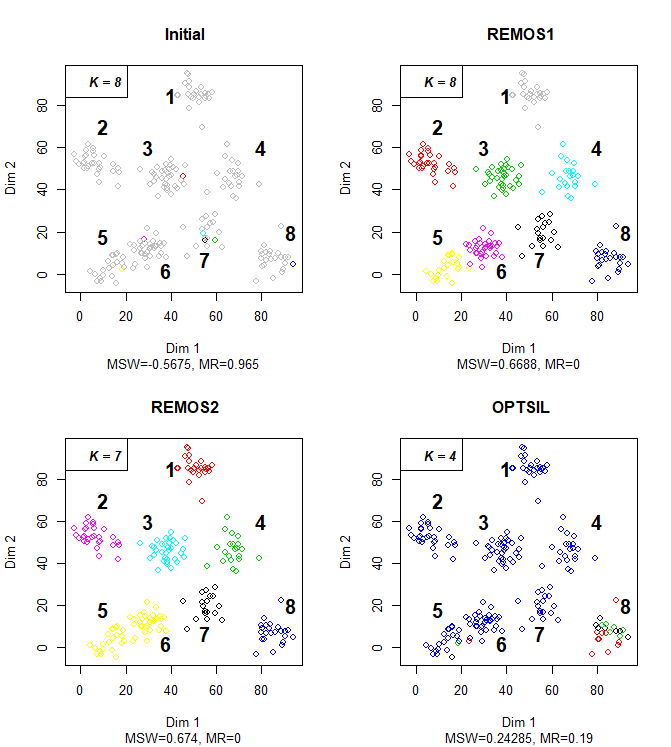
