## Supplementary material for "Comparison of silhouette-based reallocation methods for vegetation classification": Comparison of reallocation methods on real data sets using misclassification rate and the absolute sum of negative silhouette widths

Electronic Supplement for the paper

**Supplement S5 – Comparison of reallocation methods on real data sets using misclassification rate and the absolute sum of negative silhouette widths**

**Abbreviations:** MR – misclassification rate, ASNSW – absolute sum of negative silhouette widths

Figure S5-1. Comparison of the initial (without optimization) classification, REMOS1, REMOS2, and OPTSIL solutions in terms of misclassification rate and the absolute sum of negative silhouette widths across the number of clusters on the Grassland data set. The initial classification was produced by the flexible-beta method (beta = -0.25). To avoid overlap, points are jittered in horizontal direction on the graph. Colour code: red – REMOS1, green – REMOS2, blue – OPTSIL, black – initial classification.

MR and ASNSW ranked classifications very similarly.

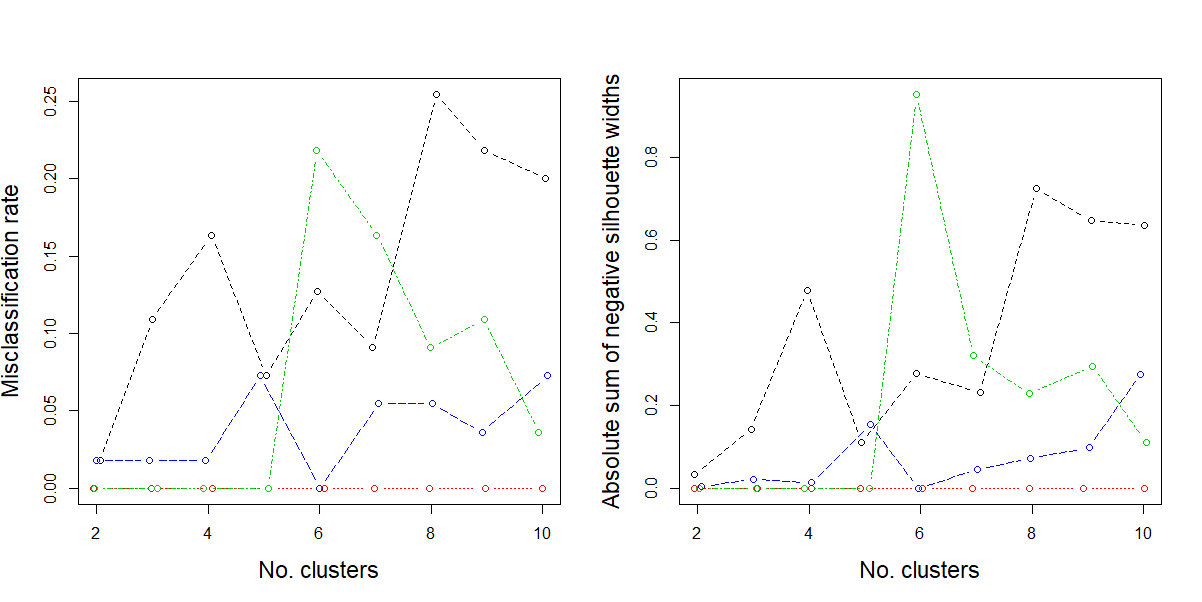

Figure S5-2. Comparison of the initial (without optimization) classification, REMOS1, REMOS2, and OPTSIL solutions in terms of misclassification rate and the absolute sum of negative silhouette widths across the number of clusters on the Bryce data set. The initial classification was produced by the flexible-beta method (beta = -0.25). To avoid overlap, points are jittered in horizontal direction on the graph. Colour code: red – REMOS1, green – REMOS2, blue – OPTSIL, black – initial classification.

Again, MR and ASNSW ranked classifications very similarly. However, OPTSIL solutions tend to seems less efficient or even worse than the initial classification from six clusters and higher.

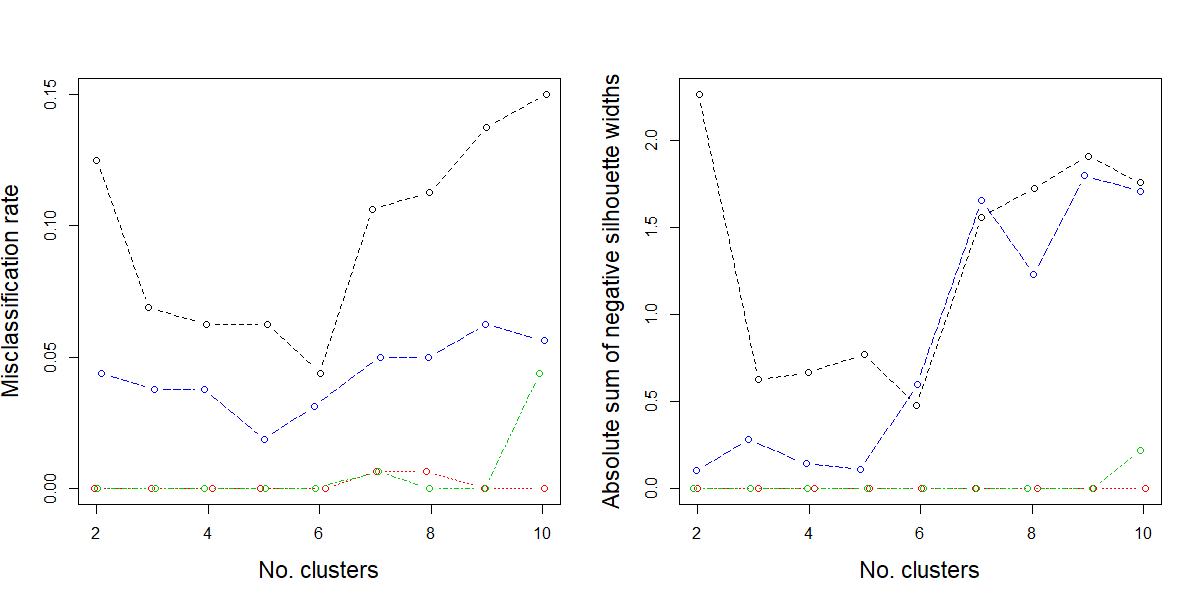

Figure S5-3. Comparison of the initial (without optimization) classification, REMOS1, REMOS2, and OPTSIL solutions in terms of misclassification rate and the absolute sum of negative silhouette widths across the number of clusters on the Shoshone data set. The initial classification was produced by the flexible-beta method (beta = -0.25). To avoid overlap, points are jittered in horizontal direction on the graph. Colour code: red – REMOS1, green – REMOS2, blue – OPTSIL, black – initial classification.

MR and ASNSW ranked classifications very similarly here as well.

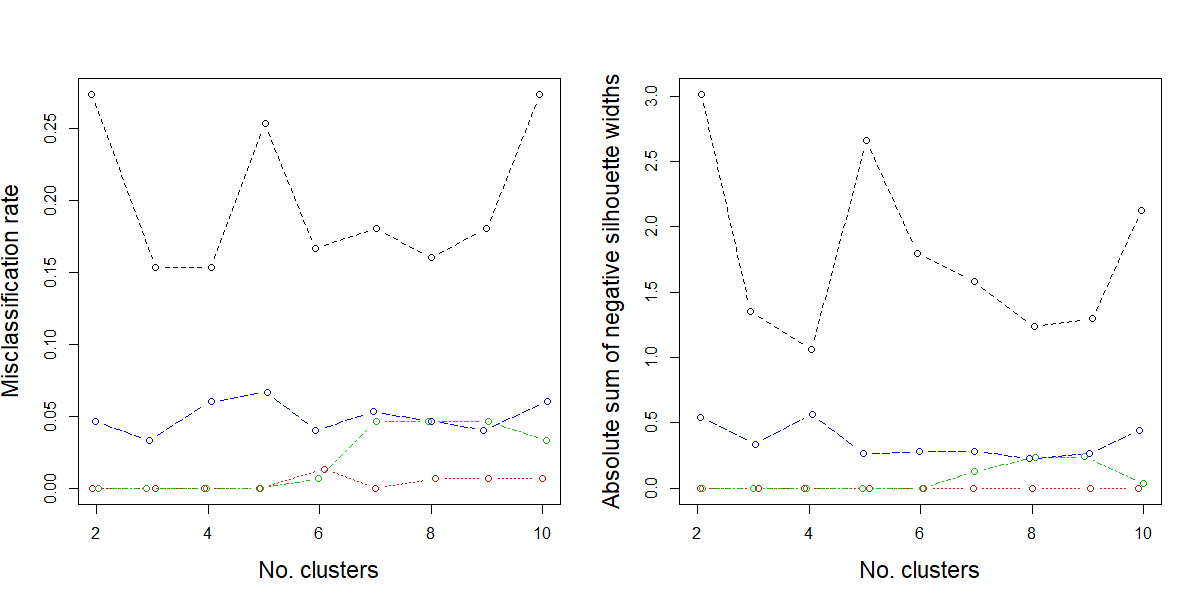
